## Supplementary material for "Trecode: a FAIR eco-system for the analysis and archiving of omics data in a combined diagnostic and research setting": Table 2

|  | **Trecode** | **OTP** | **HTS-Flow** | **QuickNGS** | **Closha** | **Terra** |
| --- | --- | --- | --- | --- | --- | --- |
| Data type | WGS, WXS, RNA-Seq, Methylation Array | WGS, WXS, WGBS, RNA-Seq, ChIP-Seq | BS-seq, RNA-Seq, ChIP-Seq, DNaseI-Seq | WGS, RNA-Seq, ChIP-Seq,miRNA | WGS,WXS, RNA-Seq, ChIP-Seq | WGS,WXS,RNA-Seq,WGBS |
| Meta data | +++ | ++ | + | ++ |  | + |
| GUI | +++ | ++ | + | + | ++ | ++ |
| Data provenance | +++ | ++ | + | + |  | + |
| Reproducibility | +++ | ++ | + | + | ++ | ++ |
| EGA/SRA compatibility | +++ | ++ | + |  |  |  |
| Automation | +++ | +++ | ++ | ++ | + | + |
| Flexibility | +++ | ++ | ++ | + | ++ | ++ |
| Workflow specification | WDL | Java/Groovy | R | bash/perl | Closha canvas | WDL |
| Workflow execution | Cromwell | Roddy/PBS/SGE/LSF | R BatchJob/SGE | Slurm, Torque | Hadoop | Cromwell |
| Execution environment | HPC Cluster, Cloud | HPC cluster | HPC cluster | HPC cluster | HPC cluster | Cloud |
| Major genomics softwares | GATK4, BWA, Samtools, STAR, MultiQC | BWA, Picard, Samtools, Platypus, ACEseq, STAR | BWA, TopHat, Bismark, MACS, Cufflinks, Cuffdiff | BWA, MACS, Samtools, Cufflinks | GATK, BWA, Samtools, STAR, TopHat, Cufflinks, Cuffdiff | GATK4, BWA, Samtools,STAR |
| Access control | * | + |  | + |  |  |
| Availability | open source | open source | open source | open source | service | service/open source** |
| Re-use | +++ | + |  |  | + | + |
| unified QC | yes (MultiQC) | no (custom) | no (custom) | no (custom) | no | no |
| Result visualization | +*** | ++ |  | ++ | + |  |
| Version control | yes | yes | yes | yes |  | yes |
| Multiuser | yes | yes | no | no | yes | yes |
| Institution | Princess Máxima Center for Pediatric Oncology | German Cancer Research Center (DKFZ) | Fondazione Istituto Italiano di Tecnologia | CECAD Research Center, University of Cologne | Korea Research Institute of Bioscience and Biotechnology | Broad Institute / Verily Life Sciences |
| Latest development | 2020 | 2020 | 2016 | 2016 | 2020 | 2020 |
|  | *=waiting for implementation in Molgenis |  |  |  |  |  |
|  | **=no documentation yet for spinning up a Terra instance | |  |  |  |  |
|  | ***=trecode includes exports to visualization platforms cBioportal and R2 | |  |  |  |  |
