## Supplementary figures and images for "Trecode: a FAIR eco-system for the analysis and archiving of omics data in a combined diagnostic and research setting"

### Supplemental Figure 1

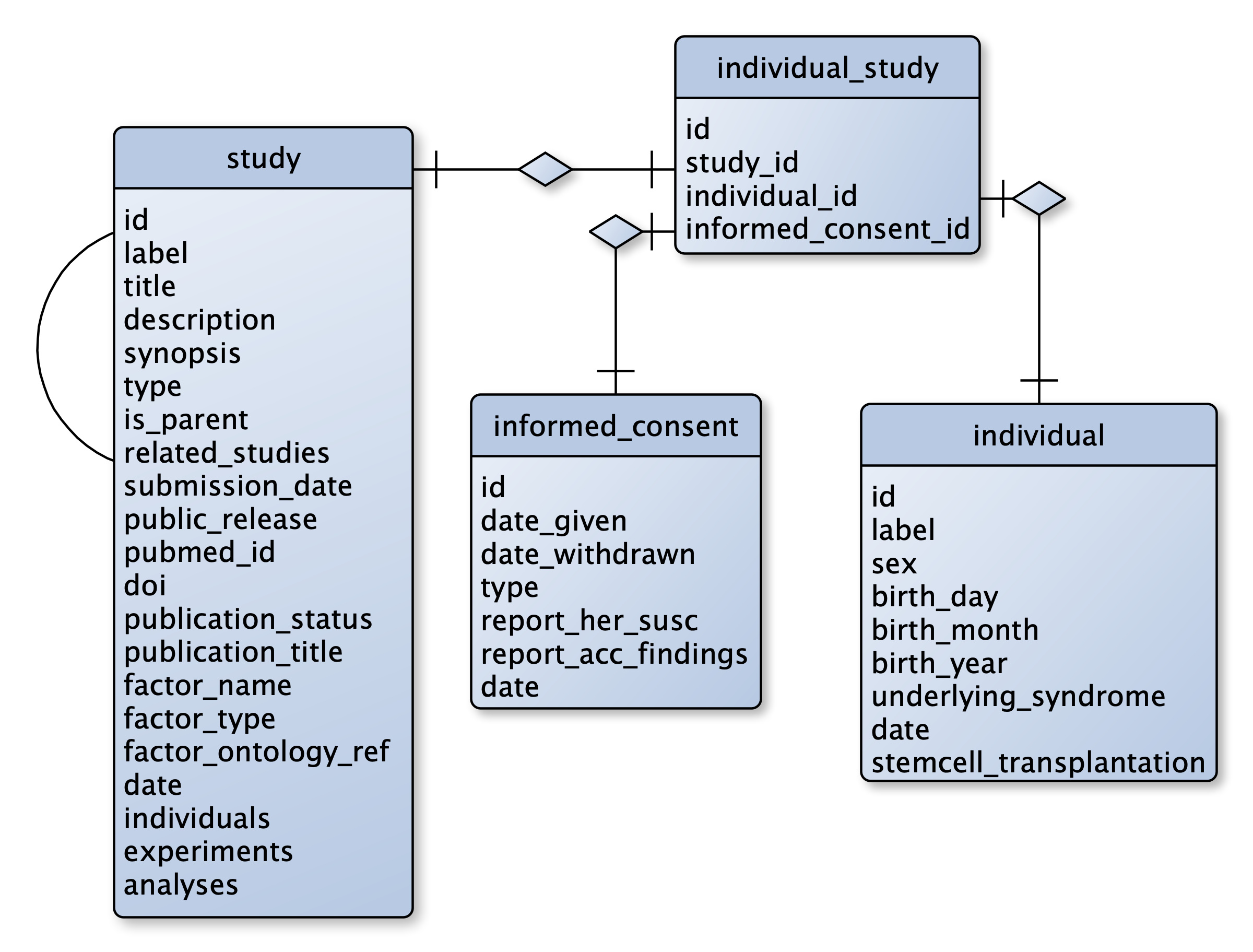

### Supplemental Figure 2

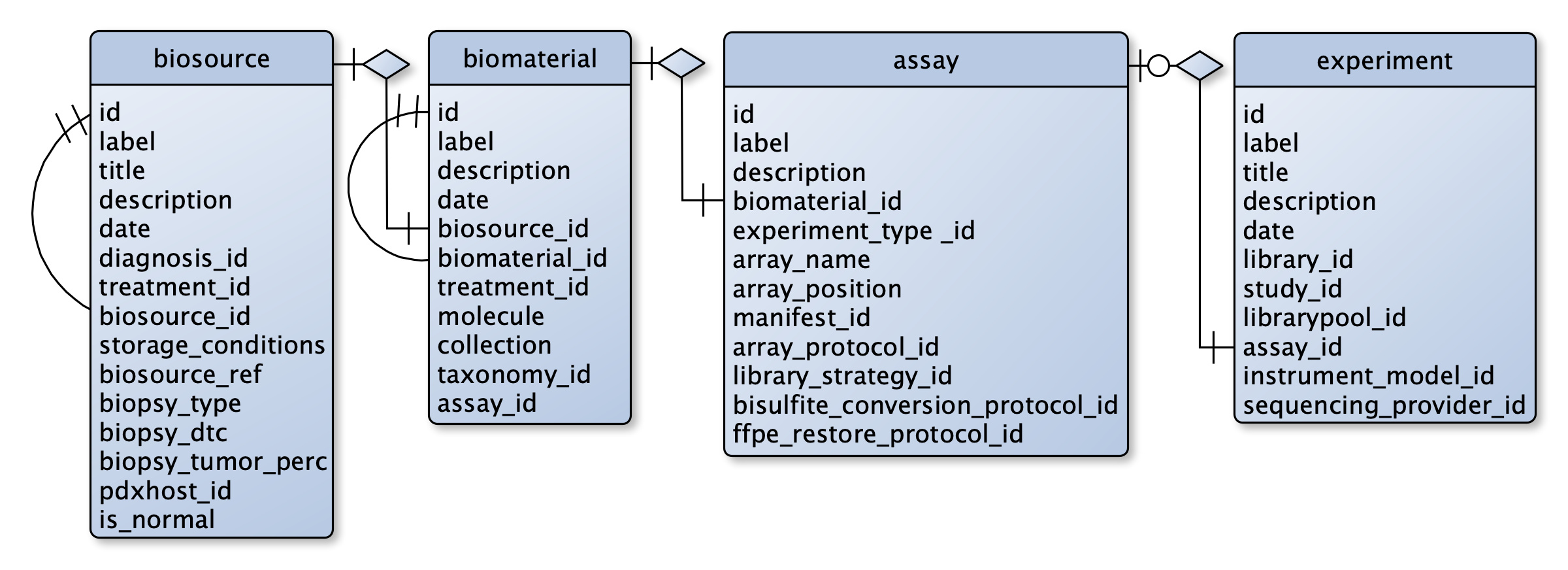

### Supplemental Figure 3a

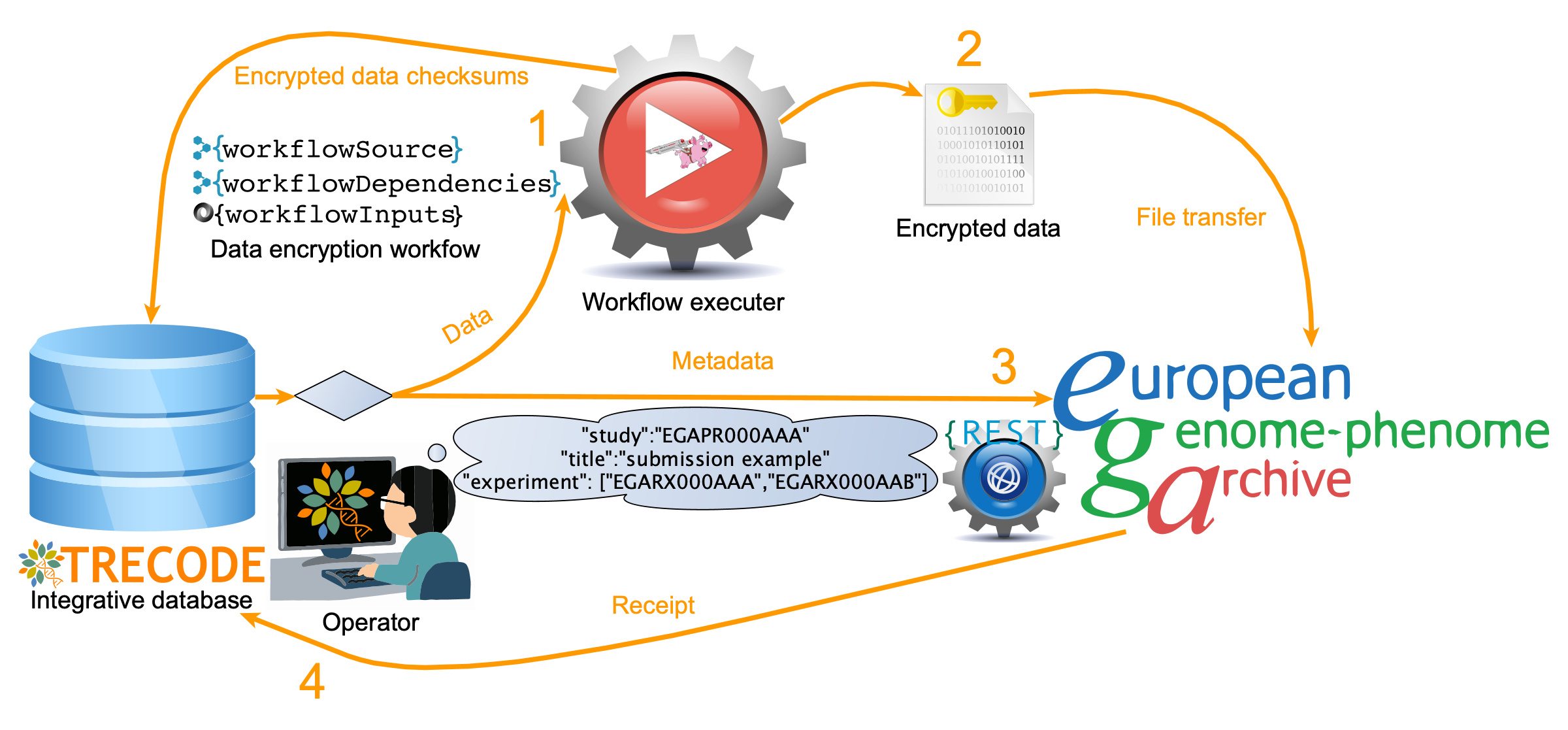

### Supplemental Figure 3b

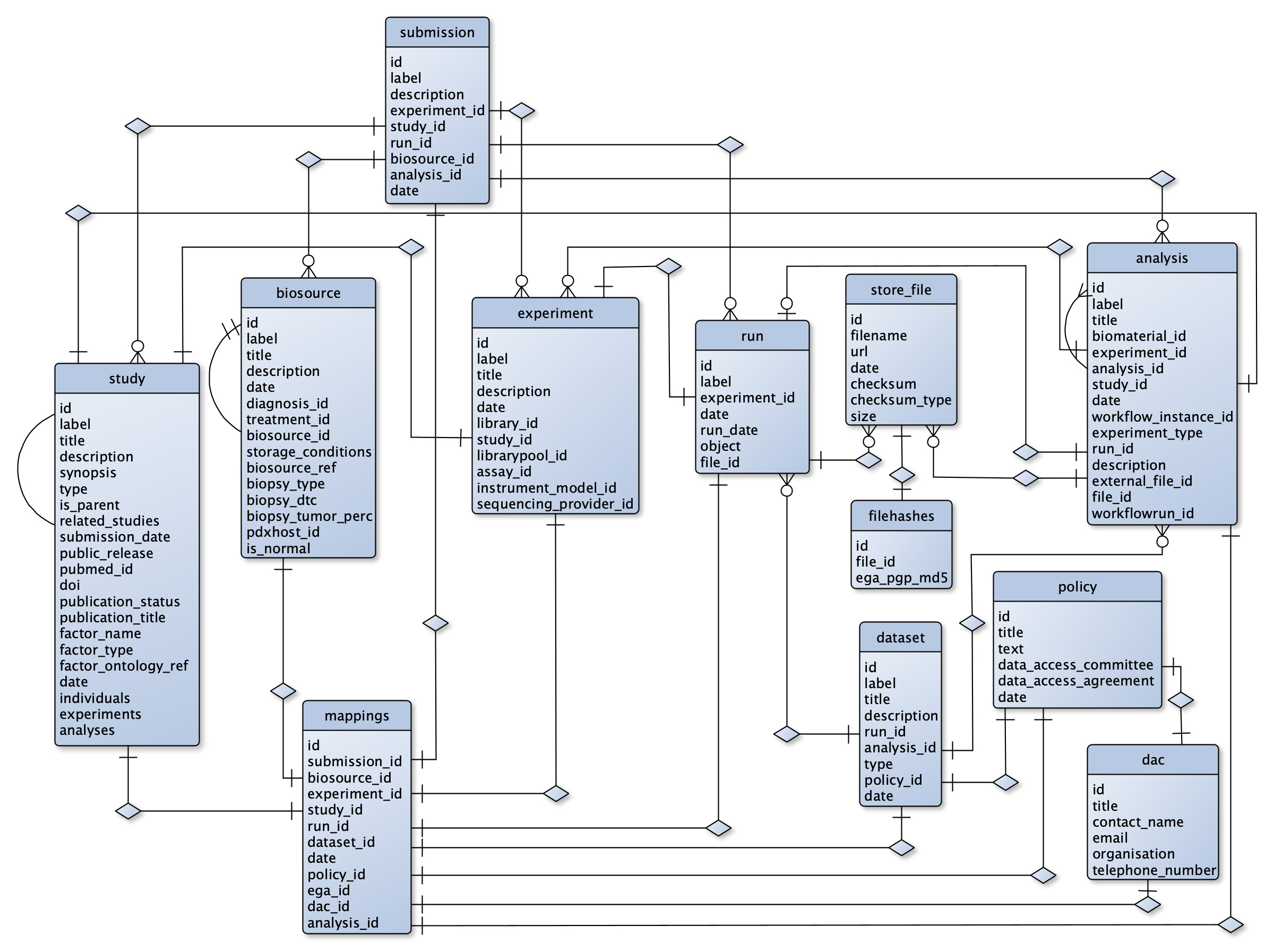
